# Targeted re-sequencing of coding DNA sequences for SNP discovery in non-model species

**DOI:** 10.1101/163659

**Authors:** Daniel W. Förster, James K. Bull, Dorina Lenz, Marijke Autenrieth, Johanna L. A. Paijmans, Robert H. S. Kraus, Carsten Nowak, Helmut Bayerl, Ralph Kühn, Alexander P. Saveljev, Magda Sindičić, Michael Hofreiter, Krzysztof Schmidt, Jörns Fickel

## Abstract

Hybridization capture coupled with high-throughput sequencing can be used to gain information about nuclear sequence variation at hundreds to thousands of loci. A cross-species approach makes use of molecular data of one species to enrich target loci in other (related) species. This is particularly valuable for non-model organisms, for which often no *a priori* knowledge exists regarding these loci. Here, we have adopted cross-species capture to obtain data for 809 nuclear coding DNA sequences (CDS) in a non-model organism, the Eurasian lynx *Lynx lynx*, using baits designed with the help of the published genome of a related model organism (the domestic cat *Felis catus*). In this manner, we were able to survey intraspecific variation at hundreds of nuclear loci across the European range of *L. lynx*. A large set of bi-allelic candidate SNPs was then tested in a high throughput SNP-genotyping platform (Fluidigm), which we reduced to a final 96 SNP-panel based on assay performance and reliability; validation was carried out with additional samples not included in the SNP discovery phase. The 96 SNP-panel developed from CDS performed very successfully in the identification of individuals and in population genetic structure inference (incl. the assignment of individuals to their source population). In keeping with recent studies, our results show that genic SNPs can be valuable for genetic monitoring of wildlife species.

## INTRODUCTION

For studies that require sequence information of only particular parts of a genome, it has become common practice to query nuclear genomes (usually of humans and model-organisms) for genetic variation using ‘enrichment’ techniques (also called ‘hybridization capture’). In this approach, so-called ‘baits’ (DNA/RNA molecules) with sequences complementary to genomic regions of interest are used to capture said regions in genetic libraries prior to sequencing, while unwanted ‘off-target’ DNA is washed away (Gasc et al., 2016). This method is so successful that commercial suppliers already offer enrichment assays for a plethora of applications (e.g. targeting loci associated with particular traits or diseases) and also offer to design custom assays. Even automation of the entire process (library building and subsequent enrichment) is already available.

For molecular, ecological and taxonomic research in non-model organisms, this technique has proven to be highly valuable. It has strongly facilitated the study of intraspecific variation of whole mitochondrial genomes, particularly when DNA from sample material is scarce and/or degraded or when the resolution of PCR-amplified single short mitochondrial sequences is insufficient to answer particular research questions (Paijmans et al., 2013). The taxonomically wide accessibility of mtDNA reference sequences for bait design together with the availability of do-it-yourself protocols for bait generation (Maricic et al., 2010) have also facilitated both its implementation and proliferation. Besides its success in mtDNA research, the technique has also been applied to nuclear loci in non-model organisms, but here mainly in interspecific studies to address taxonomic uncertainties. The promiscuity of the baits in the hybridization capture process allows for up to 40% sequence divergence between bait and target (Li et al., 2013), a property that can be exploited to obtain relatively complete datasets of 100s-1000s loci (Lemmon et al., 2012; Faircloth et al., 2012; Bi et al., 2012; Prum et al., 2015) across different taxonomic levels (genera, families, even orders).

In this study, we demonstrate the application of hybridization capture for single-nucleotide polymorphism (SNP) discovery in a non-model species and show how the identified SNPs can be utilized for cost-effective genetic monitoring of an elusive carnivore, the Eurasian lynx *Lynx lynx*.

We used a ‘cross-species capture’ approach to enrich target loci from our study species (lynx) using baits designed from a fully annotated reference genome of a related species, namely the domestic cat. In order to avoid capturing paralogues, yet have loci evenly distributed throughout the lynx genome, we designed cat baits to target single-copy coding DNA sequences (CDS). To minimize ascertainment bias, lynx samples used for SNP discovery covered the European distribution range of the species, including some reintroduced populations.

In the past, the applicability of SNPs for monitoring wildlife populations was quite limited, because the (usually very low) amount of template DNA extractable from non-invasively collected samples constrained the number of SNP loci that could be genotyped in such samples (e.g. Kraus et al., 2015, and references therein). However, technological advancements such as nanofluidics substantially reduced the required reaction volumes for SNP genotyping, making it possible to simultaneously type numerous SNP loci even from very little template material. For this reason, we aimed to develop a SNP-panel that can be routinely used with the nanofluidic Dynamic Array Chip technology implemented in the Fluidigm platform (Fluidigm Corp., SanFrancisco, CA, USA).

Here, we report the development of a 96 SNP-panel for the Eurasian lynx *Lynx lynx* (Fig. 1A). We outline how we (i) used the genomic resources available for model organisms to design baits and then enriched target loci in our study species; (ii) filtered the intraspecific variation in our study species for candidate SNP loci; (iii) evaluated a large set of candidate SNPs in our chosen genotyping platform, and (iv) settled on 96 loci for the final SNP-panel. Then, we present how the newly developed SNP-panel performed in the genetic monitoring of our study species using additional samples (not included in the SNP discovery). To evaluate performance, we specifically focused on the ability of a SNP-panel developed from coding DNA sequences for (i) individual discrimination and (ii) analysis of genetic population structure, including the correct assignment of individuals to their source population.

**Figure 1.**
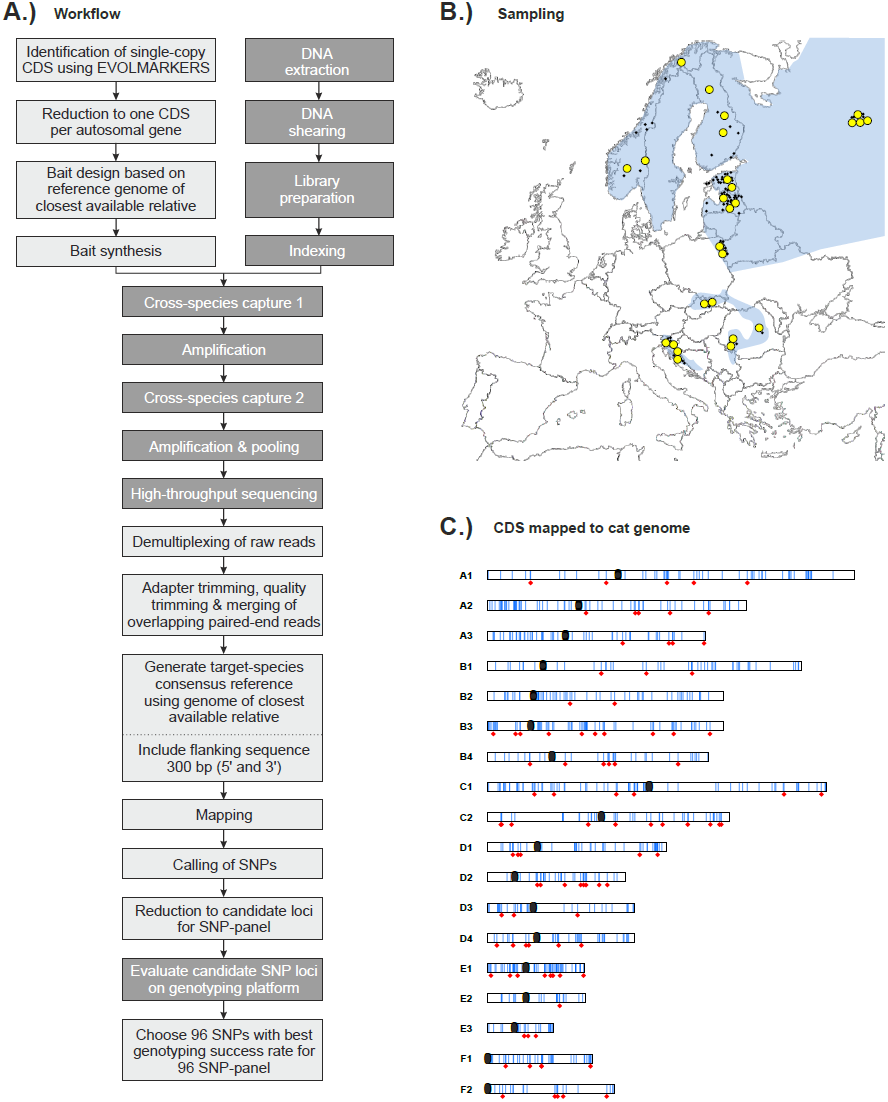
Schematic overview of study. (A) Schematic summary of the workflow, presenting both computational steps (light grey boxes) and laboratory steps (dark grey boxes). (B) Sampling localities of Eurasian lynx *Lynx lynx* across Europe. The current distribution of the species is shaded in blue. Large yellow circles represent samples used during the SNP discovery phase; small black dots represent additional samples genotyped using the developed 96 SNP-panel. (C) Schematic representation of the distribution of CDS targeted for enrichment (blue bars), projected onto cat chromosomes. The positions of the 96 SNP loci used for the SNP-panel are indicated by red diamonds. Black ovals show centromeres.

## MATERIALS AND METHODS

### Biological samples

DNA was extracted from tissue (liver or muscle) or blood using the commercial First-DNA all tissue kit (GEN-IAL GmbH, Troisdorf, Germany). 26 Eurasian lynx from four European populations (von Arx et al. 2004) were used in the initial cross-species capture: *Baltic –* Estonia (*N* = 2), Latvia (*N* = 3), Poland (*N* = 2), Russia (*N* = 4); *Nordic –* Finland (*N* = 3), Norway (*N* = 3); *Carpathian –* Romania (*N* = 3), Slovakia (*N* = 2); and *Dinaric –* Croatia (*N* = 2), Slovenia (*N* = 2) (Table 1, Fig. 1B).

**Table 1.** Sampling localities, sequencing results and cross-species capture results of 26 Eurasian lynx *Lynx lynx.*

| Sample ID | Country | Population <sup>a</sup> | Raw reads | Filtered sequences <sup>b</sup> | On-target reads (%) | % of target covered at 15x <sup>c</sup> |
| --- | --- | --- | --- | --- | --- | --- |
| Cro.1 * | Croatia | reintroduced | 42 321 520 | 28 717 105 | 19.6 | 79.0 |
| Cro.2 | Croatia | reintroduced | 19 467 436 | 15 178 879 | 26.2 | 66.2 |
| Est.1 | Estonia | natural | 20 936 570 | 15 527 441 | 24.6 | 80.3 |
| Est.2 | Estonia | natural | 25 697 066 | 17 894 280 | 25.6 | 81.9 |
| Fin.1 | Finland | natural | 22 477 300 | 13 883 204 | 27.6 | 82.4 |
| Fin.2 | Finland | natural | 22 967 626 | 16 569 199 | 24.7 | 82.8 |
| Fin.3 | Finland | natural | 22 894 258 | 16 450 208 | 24.2 | 63.4 |
| Latv.1 | Latvia | natural | 22 321 270 | 17 341 768 | 23.6 | 77.7 |
| Latv.2 | Latvia | natural | 24 394 838 | 18 562 462 | 23.7 | 80.1 |
| Latv.3 | Latvia | natural | 21 356 226 | 16 677 724 | 20.8 | 81.6 |
| Nor.1 | Norway | natural | 25 157 782 | 15 829 112 | 26.5 | 66.3 |
| Nor.2 | Norway | natural | 25 893 728 | 18 727 706 | 24.1 | 82.9 |
| Nor.3 * | Norway | natural | 51 194 776 | 33 117 564 | 19.9 | 81.8 |
| Pol.1 | Poland | natural | 36 350 048 | 22 862 227 | 27.0 | 83.1 |
| Pol.2 | Poland | natural | 23 139 694 | 15 996 340 | 26.8 | 73.0 |
| Rom.1 | Romania | natural | 23 078 140 | 17 044 276 | 27.4 | 83.1 |
| Rom.2 | Romania | natural | 24 678 618 | 16 330 922 | 29.5 | 83.2 |
| Rom.3 * | Romania | natural | 45 479 818 | 28 824 269 | 21.3 | 78.3 |
| Rus.1 | Russia | natural | 28 224 284 | 20 381 841 | 22.9 | 80.6 |
| Rus.2 * | Russia | natural | 36 384 372 | 25 422 519 | 19.9 | 83.0 |
| Rus.3 | Russia | natural | 24 140 510 | 18 351 015 | 23.1 | 81.0 |
| Rus.4 | Russia | natural | 23 737 994 | 17 874 068 | 22.8 | 81.6 |
| Svk.1 | Slovakia | natural | 21 955 584 | 16 006 621 | 29.2 | 82.8 |
| Svk.2 | Slovakia | natural | 25 109 422 | 18 732 792 | 26.8 | 83.7 |
| Svn.1 | Slovenia | reintroduced | 24 523 316 | 18 384 101 | 24.9 | 85.1 |
| Svn.2 * | Slovenia | reintroduced | 46 579 692 | 28 828 601 | 23.0 | 86.8 |
| *mean* |  |  | 28 094 688 | 19 596 779 | 24.4 | 76.7 |

\* incl. data from single capture (see METHODS).
<sup>a</sup> ‘natural’ refers to autochthonous populations or populations that have recovered from population bottlenecks through natural means (i.e. without human assistance); ‘reintroduced’ populations are those established through human assistance (translocations, etc).
<sup>b</sup> adapter and quality trimmed, merged sequences and sequences that could not be merged
<sup>c</sup> of the targeted 809 CDS with a total length of 618 547 bp (i.e. not incl. flanking sequences).

Applicability of the final 96 SNP-panel was then assessed by genotyping an additional 96 lynx samples originating from the same populations (below, but see also supplementary Table S1).

The *Dinaric* population was intentionally sampled as a distinct reintroduced population (originating from the Carpathian population), for the assessment of correct individual assignment (below).

### Bait design

We compared the annotated genomes of the domestic cat (*Felis catus* v6.2), domestic dog (*Canis lupus familiaris* v3.1), horse (*Equus caballus* v2.0), cow (*Bos taurus* v3.1) and pig (*Sus scofa* v10.2) using EVOLMARKERS (Li et al., 2012) to identify single-copy protein coding genes present in all of these taxa. In order to avoid paralogues, candidate target loci were restricted to CDS with less than 40% similarity to intraspecific sequences using a BLAST approach (Li et al., 2012). This restriction ensured that target loci would be unambiguously identifiable. To increase the chance of SNP discovery and facilitate the development of assays, we excluded short sequences and thus chose targets with a minimum length of 400 bp, which yielded 1357 CDS markers. Potential targets were then further filtered by selecting a single CDS per autosomal gene, reducing the set to 809 CDS markers. As capture was to be performed on a felid species, we used *Felis catus* CDS to design the baits (Fig. 1C, see also information summary in supplemental Table S2). The custom tailored MYbaits^®^ target enrichment kit (MYcroarray, Ann Arbor, MI, USA) covering all 809 CDS (having a total length of 618 547 bp) finally consisted of 8922 biotinylated RNA baits (2× tiling, 120 bp length). The bait design is available as a FASTA file on DRYAD (doi: provided upon acceptance).

### Capture of CDS and sequencing

Illumina sequencing libraries were built following a published DIY protocol (Meyer & Kircher, 2010) with some modifications reducing both loss of template and costs (Fortes & Paijmans, 2015).

Libraries were captured individually following suggested modification of the MYbaits protocol (Li et al., 2013). In brief, volumes were reduced, the amount of synthetic RNA-baits used per capture was reduced (8-fold), and the standard hybridization temperature of 65 ° C was replaced by a ‘touchdown’ protocol with hybridization temperature decreasing from 65 ° C to 50 ° C in 5 ° C increments every 11 hrs. Detailed methods for library building and capture can be found in the Appendix.

The library of each individual (*N* = 26) was captured two consecutive times; the eluate from the first capture (i.e. the enriched library) was amplified and used in a second round of capture, as this has been reported to increase the number of on-target reads for both within-species and cross-species capture (Li et al., 2013; Templeton et al., 2013). We confirmed this by sequencing the first five samples processed also after the first capture round, observing a four-to seven-fold increase in the number of unique on-target sequences after two consecutive captures (supplemental Fig. S1).

Enriched libraries were paired-end sequenced on two Illumina platforms (Illumina, San Diego, CA, USA): MiSeq using v2 300-cycle kits, and NextSeq using v1 150-cycle MidOutput kits.

### Data processing

#### Pre-Processing

De-multiplexing of paired-end reads using BCL2FASTQ v2.17.1.14 (Illumina, Inc.) was followed by removal of adapter sequences using CUTADAPT v1.3 (Martin, 2011). Adapter-clipped reads were then quality-trimmed using a sliding window approach in TRIMMOMATIC (Bolger et al., 2014), with the phred quality threshold set at Q=20 and window length of 10bp. Overlapping paired-end reads having a minimal overlap of 10 bp were merged using the software FLASH v1.2.8 (Magoč & Salzberg, 2011). Merged and non-merged sequences were used as input for mapping using BURROWS-WHEELER ALIGNER V0.7.10 (BWA; Li & Durbin, 2009) with seeding disabled (Schubert et al., 2012).

#### Reference processing

Because lynx CDS display little sequence divergence from orthologous cat CDS (~0% - 4% sequence divergence; DWF, unpublished data), we were able to use a simple approach to generate lynx CDS references. First, we retrieved 300bp of flanking sequences (both 5’ and 3’) of each cat CDS (cat genome v8.0) to extend the cat reference sequence beyond CDS boundaries. This was done by querying the CDS bait sequences vs. the *Felis catus* v8.0 genome using BLASTN (BLAST+ v2.2.29); the resulting hits were restricted to only one target hit (-max_target_seqs 1) and were further filtered to match only the predicted chromosome. The actual CDS sequence including flanking sequence was retrieved by applying BEDTOOLS *getfasta* (v2.17.0) using ±300bp CDS positions on the cat reference. Then, these extended cat sequences (CDS + flanks) served as reference for mapping all lynx sequences (i.e. from all samples). Aligned sequences were de-duplicated using *MarkDuplicates* from PICARD-TOOLS v1.106 (https://github.com/broadinstitute/picard). Variant calling was carried out using SAMTOOLS v1.1 (Li et al., 2009) and BCFTOOLS v1.2 (http://github.com/samtools/bcftools) to determine the most common lynx variant at every position; the cat reference was modified accordingly. For variants present at ≥3× and alternate base frequency > 0.5 the alternate (lynx) base was used to generate the lynx consensus sequence. In this manner, we converted the cat sequence into a lynx ‘consensus’ sequence for each CDS, which included 300 bp of 5’ and 3’ flanking sequence; henceforth, the ‘lynx CDS reference’ (of 1,102,167 bp length). We carried out the same procedure using the recently published Iberian lynx (*Lynx pardinus*) genome as reference (Abascal et al., 2016), representing a closer relative of our study species, albeit with a less complete genome assembly. It should be noted, however, that in cases where the target species is highly divergent from the reference species, other approaches are advised to generate target species reference sequences (e.g. Yuan et al., 2016; Portik et al., 2016).

#### Sample Processing

We generated a separate consensus sequence for each lynx sample by mapping against the lynx CDS reference. Following a second round of mapping against this sample-specific consensus sequence (in order to recover as much data as possible), GATK *UnifiedGenotyper* (v1.6) was then used to identify sequence variants (SNPs and InDels) in the 26 Eurasian lynx samples used for SNP discovery. Of all SNPs identified, only SNPs with a coverage ≥15× were retained as candidates for the SNP panel. These candidate loci were then reduced to 144 SNPs (representing a 50% surplus over the target number of 96 SNPs for the final panel) based on the following criteria: (i) SNPs had to be bi-allelic (a requirement of the genotyping platform); (ii) minor allele frequency should be greater than 10% (to avoid loci with rare alleles and hence potentially low applicability); (iii) no variants (SNPs or InDels) within 100 bp of the candidate SNP to avoid interference in the genotyping process; (iv) only one SNP allowed per CDS to avoid physical linkage; (v) SNPs should be as widely distributed across the genome as possible (using the cat genome as reference, Fig. 1c); and (vi) SNPs should not lay in the flanking regions of the CDS, because our goal was to create a SNP panel from data obtained following cross-species enrichment of CDS (flanking sequences may not be available for all study species).

### SNP-panel development

We aimed to generate a lynx SNP-panel with 96 SNPs for high-throughput genotyping using Fluidigm’s SNPtype™ assays (Fluidigm Corp., SanFrancisco, CA, USA). Specifically, we intended to use the ‘96.96 Dynamic Array Chip for Genotyping’ that allows the simultaneous genotyping of 96 samples at 96 bi-allelic SNPs, which is particularly useful when only little sample material is available. The latter was an important criterion for platform selection because non-invasively collected samples represent an important resource for genetic monitoring of wildlife populations, and these often yield very little DNA for analysis, which thus needs to be used very efficiently. The nanofluidic Dynamic Array Chip technology employed in this platform reduces PCR volumes to nanolitres, performing 9216 (96×96) single-plex reactions in a highly automated fashion. As little as 1.25 μL of DNA extract (0.5 ng/μL) is sufficient to genotype one sample at 96 SNPs. Genotyping itself is accomplished using allele-specific primers labeled with fluorescent dyes. For samples with low amounts of template (e.g. non-invasively collected samples) a pre-amplification is strongly recommended (Nussberger et al., 2014; Kraus et al., 2015).

Prior to the selection of 96 SNPs for the final genotyping panel we pre-selected 144 candidate SNPs (96 + 48) for evaluation (above), of which we randomly chose 15 (10.4%) for verification in five of the 26 Eurasian lynx samples using Sanger sequencing. After their successful verification, we designed SNPtype™ assays for all 144 SNPs.

Using previously established procedures (Kraus et al., 2015) we assessed genotyping errors for all 144 SNPtype™ assays. Specifically, we genotyped the 26 Eurasian lynx samples that had been sequenced for SNP discovery using both undiluted (50 ng/μL) and diluted (0.5 ng/μL) samples. The latter served to approximate poor DNA quality samples (e.g. coming from non-invasively collected samples). We used ‘genotyping treatment c2’ (Kraus et al., 2015; less dilution of specific target amplification [STA] products, 42 cycles of amplification), and evaluated (by locus) the following properties: genotype consistency across dilutions, the incidences of missing data, and genotype consistency with the Illumina sequencing data.

For the assembly of the final 96 SNP-panel, we chose the assays with the most consistent genotyping performance across the 26 lynx samples (i.e. showed lowest rates of missing data and genotyping errors) and for which the three genotypes (AA, AB, BB) could be unambiguously distinguished in the scatter plots generated as output by the EP1 genotyping software (Fluidigm Corp.). To test the general applicability of the final 96 SNP-panel, we genotyped a set of 96 Eurasian lynx samples from across Europe (Table S1), none of which had been used in the SNP discovery. Again, we randomly chose 10 out of 96 loci (10.4%) to be verified by Sanger sequencing in five out of the 96 Eurasian lynx.

As DNA of prey species may contaminate DNA extracted from non-invasively collected fecal samples, we also performed cross-species testing of the SNP-panel on typical representatives of common prey taxa: roe deer *Capreolus capreolus*, European red deer *Cervus elaphus*, European hare *Lepus europaeus*, house mouse *Mus musculus*, pine marten *Martes martes*, and American mink *Neovison vison*.

### Individual identification

We used two approaches to examine the power of the newly designed 96 SNP-panel to discriminate among individuals. Firstly, we used GIMLET v.1.3.3 (Valière, 2002) to estimate (across loci) both the unbiased ‘probability of identity’ (PID_unb_) and the more conservative ‘probability of identity given siblings’ (PID_sib_). These probabilities were estimated separately for each population with at least 10 individuals (Estonia, Latvia, Poland, Norway, Slovenia), as well as for the genetic clusters identified in the population structure analyses (below).

Secondly, we directly examined the performance of the SNPs to differentiate all individuals in our data set. Specifically, we examined how well different subsets of loci performed for identifying individuals. We examined a range of subset sizes (10, 20, 30, 40, 50, 60, 70, 80, and 90 loci), and examined the results for 10,000 permutations per subset size; for this, subsets of loci were randomly drawn from the entire dataset without replacement. This analysis was conducted in the statistical programming environment R (http://www.cran.r-project.org) using a custom script (deposited on DRYAD under doi: provided upon acceptance).

### Population structure inferences

For the assessment of SNP performance regarding the detection of genetic substructure and population assignment we used two common methods: principal component analysis (PCA) and Bayesian population assignment. PCA was carried out using the R package *adegenet* v.2.0.1 (Jombart, 2008). Bayesian assignment to genetic clusters (populations) was carried out using the software STRUCTURE v2.3 (Pritchard et al., 2000). In the latter, we were interested in the number of genetic clusters identified, but also if the SNPs could be used to assign lynx to their correct source cluster. To examine this, we first conducted the STRUCTURE analysis on samples (*N* = 99) from naturally occurring populations and then tested whether samples from reintroduced populations (Croatia, Slovenia; *N =* 20) would be assigned to the genetic cluster corresponding to their source population, i.e. the population from which lynx had been translocated to establish the reintroduced populations. First, for all 99 lynx from naturally occurring populations, we ran 10 replicates for values of *K* (inferred number of genetic clusters) from 1 to 8 for 600,000 iteration steps, the first 150,000 of which were discarded as burn-in, and allowing for correlated allele frequencies in the admixture model (Falush et al., 2003). The most likely number of genetic clusters (*K*) was then determined following the Δ*K*-method (Evanno et al., 2005) implemented on the STRUCTUREHARVESTER website (http://taylor0.biology.ucla.edu/structureHarvester) (Earl & vonHoldt, 2012). Using the inferred *K*, we then re-ran STRUCTURE with all lynx (*N* = 119) to check for correct cluster assignment.

We tested for linkage disequilibrium (LD) and deviation from Hardy-Weinberg equilibrium (HWE) with GENEPOP (Raymond & Rousset, 1995) in sampling localities and inferred genetic clusters; the R package *LDheatmap* (Shin et al., 2006) was used to visualize pairwise *r*^2^ values.

### Applicability of baits for other taxa

To assess the potential taxonomic breadth for which our cat CDS derived baits may be applicable, we queried our baits against the published genomes of other carnivore species using BLASTN: Iberian lynx (*Lynx pardinus* v1.0), cheetah (*Acinonyx jubatus* v1.0), leopard (*Panthera pardus* v1.0), tiger (*Panthera tigris altaica* v1.0), dog (*Canis lupus familiaris* v3.1), ferret (*Mustela putorius furo* v1.0), polar bear (*Ursus maritimus* v1.0), and giant panda (*Ailuropoda melanoleuca* v1.0). As sequence divergence between bait and target impacts enrichment (Vallender, 2011; Bi et al., 2012; Hedtke et al., 2013; Peñalba et al., 2014; Paijmans et al., 2016), we determined the sequence similarity between the baits and their corresponding best resulting hit in the queried genomes.

## RESULTS

### Sequence and target-enrichment results

The number of raw reads varied from 19,467,436 to 46,579,692 among the 26 lynx samples (mean: 28,094,688), and the percentage of on-target sequences ranged from 19.6% to 29.5% (mean: 24.4%; Table 1). As the target region corresponds to 0.026% of the cat genome (supplemental Table S2), and the lynx genome is roughly equivalent in size (Animal Genome Size Database, http://www.genomesize.com), the cross-species capture resulted in a greater than 900-fold increase in on-target sequences.

For half of the CDS the enrichment was very successful: on average, 90% or more of the target region was covered at ≥15× depth (Fig. 2A). Across all samples, the majority of CDS (632 loci; 78.1%) had a coverage of ≥15× for 50% or more of their lengths. Some CDS (61 loci; 7.5%) were poorly enriched in all samples (less than 10% of target region ≥15× depth), among them 31 CDS were not enriched in any sample.

**Figure 2.**
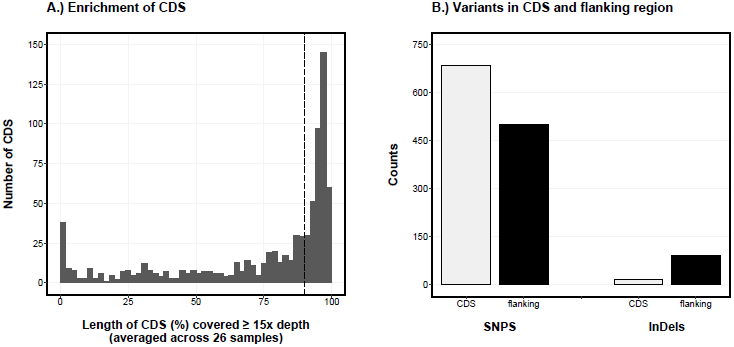
(A) Average recovery of the 809 CDS enriched in the 26 Eurasian lynx in the SNP discovery panel. For each CDS, the average percentage of the locus length covered at ≥15× depth is plotted (using 2% bins). 50% of CDS have above 90% sequence length coverage at ≥15× depth, indicated by the black vertical line. (B) Summary of the distribution of variation in CDS and flanking sequences, shown separately for SNPs and InDels.

As expected, the inclusion of flanking sequences in the lynx CDS references (300 bp, both 5’ and 3’ see METHODS) increased the number of mapped sequences and improved coverage at CDS boundaries. This yielded between 16,701 and 30,309 more bases of the target with ≥15× depth (mean: 20,674 bases; 3.34% of the target region).

### Variation in CDS

Of the 809 CDS analyzed, 61 were excluded due to insufficient data, 207 showed no variation among lynx samples (most probably due to poor coverage of CDS at ≥15× depth) and one showed no variation between lynx and cat. Three further CDS were no longer present in the newest build of the cat genome (v8.0). The remaining 537 CDS (66.4% of CDS; length ranging from 961 to 5113 bp incl. flanking sequence) showed intraspecific variation, which consisted of 1186 SNPs and 109 InDels (Fig. 2B).

Use of the Iberian lynx genome as reference (see METHODS) revealed fewer intraspecific variants among Eurasian lynx samples (993 SNPs and 66 InDels). Most of this difference in the number of variants detected reflects differences in the genome assemblies of the cat and the Iberian lynx (i.e. fewer CDS present in the genome assembly, less flanking sequence retrievable) – in other words, the variation detected using the Iberian lynx genome was mostly a subset of the variation detected using the cat genome. Variants detected only when using the Iberian lynx genome as reference were located in the flanking regions of 14 CDS (18 SNPs, 3 InDels), which were characterized by low or no coverage using the cat genome. Thus, although the Iberian lynx is a closer relative of the Eurasian lynx than is the domestic cat, the more complete genome assembly of the cat enabled us to retrieve more data about intraspecific variation in the Eurasian lynx.

### SNP-panel

Out of the 686 candidate SNPs inside CDS (57.8% of SNPs), we selected 144 for further evaluation (see METHODS for selection criteria). This number was then further reduced to 96 SNPs based on reliability for genotyping on the chosen platform (Fluidigm Corp., ‘96.96 Dynamic Array Chip for Genotyping’). Based on the distribution of the selected SNPs when projected onto the cat genome (Fig. 1C) and assuming a similar distribution in lynx, we estimated the average distance between SNPs on the same chromosome in the final set of 96 SNPs to be 17.55 Mb.

As indicated by the improved genotyping success (Fig. 3), the final set of 96 SNPs had less missing data or genotyping inconsistencies among sample replicates of the 26 lynx, than did the set of all 144 SNPs. Five out of the 26 diluted samples (0.5 ng/μL of DNA), which approximated poor quality samples (e.g. non-invasively collected samples), had a high incidence of missing data (visible as outliers in Fig. 3 left and mid). We obtained complete or nearly complete genotypes for 93 of the 96 additional lynx samples (Fig. 3 right, mean genotyping success = 97%). We found no significant linkage between loci (LD heatmap in suppl. Figure S2), and detected no deviation from Hardy-Weinberg equilibrium in sampling localities or inferred genetic clusters (below).

**Figure 3.**
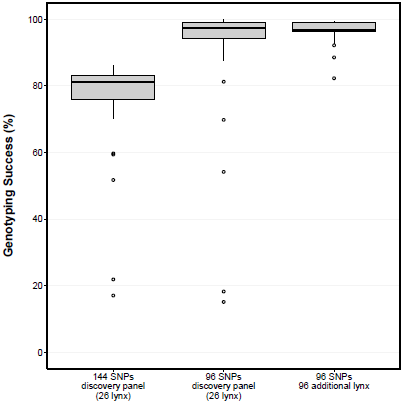
Box plots displaying the genotyping success of Eurasian lynx samples at SNP loci: left, the discovery panel (26 lynx) for the initial set of 144 SNPs tested on the Fluidigm genotyping platform; middle, the final 96 SNP-panel genotyped on the discovery panel (26 lynx); and right, the final 96 SNP-panel on an additional 96 lynx (not part of discovery panel).

Cross-species tests resulted in low overall genotyping success, except for the domestic cat (upon which the cross-species capture was based), which generated data at 83 loci, out of which only five were heterozygous (6%). The other prey species displayed signals at a far lower rate: roe deer at 13 loci, European red deer at 9 loci, European hare at 8 loci, house mouse at 2 loci, pine marten at 27 loci, and American mink at 24 loci. Thus, the most likely prey species (roe deer, red deer, hare, [Jobin et al., 2000; Belotti et al., 2014]) would not generate false positive lynx genotypes; for the domestic cat (and presumably also the wild cat *Felis silvestris*), species-identification using commonly employed mtDNA markers may be necessary to exclude false positive lynx genotypes.

### Identification of individuals

All 96 SNPs of the final panel were polymorphic for the complete lynx dataset (*N* = 119), with a minor allele frequency (MAF) between 7% and 50% (mean = 29%). The cumulative PID_unb_ (unbiased probability of identity) within geographic localities ranged from 7.13×10^-18^ to 2.31×10^-34^ (Table 2); and in the genetic clusters identified in the population structure analyses (below) from 3.54×10^-22^ to 3.76×10^-34^. The more conservative estimate of PID_sib_ (probability of identity given siblings) ranged from 1.53×10^-8^ to 3.39×10^-17^ within geographic localities, and from 3.61×10^-11^ to 5.12×10^-17^ in genetic clusters (Table 2). As little as 24 loci were already sufficient to achieve a PID_sib_ <10^-4^, regardless of locality or genetic cluster (Table 2). When we examined the performance of SNPs to differentiate all 119 lynx using various subsets of loci (ranging from 10 to 90 loci; Fig. 4), we found that 60 SNPs were sufficient to differentiate all individuals in more than 98% of 10,000 random permutations of SNP loci (sampled without replacement). Thus, given the observed genotyping success rate (Fig. 3), our final 96 SNP-panel should perform very well for individual identification.

**Table 2.**
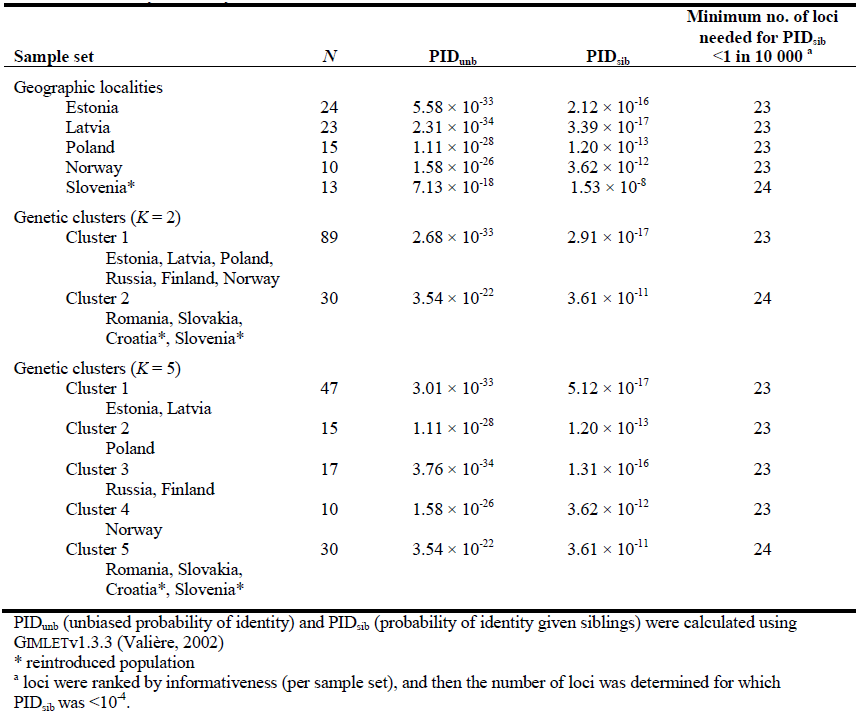
Probability of identity calculated for different subsets of the SNP data.

| Sample set | *N* | $PID_{unb}$ | $PID_{sib}$ | Minimum no. of loci<br>needed for $PID_{sib}$<br><1 in 10 000 <sup>a</sup> |
| --- | --- | --- | --- | --- |
| Geographic localities |  |  |  |  |
| Estonia | 24 | $5.58 \times 10^{-33}$ | $2.12 \times 10^{-16}$ | 23 |
| Latvia | 23 | $2.31 \times 10^{-34}$ | $3.39 \times 10^{-17}$ | 23 |
| Poland | 15 | $1.11 \times 10^{-28}$ | $1.20 \times 10^{-13}$ | 23 |
| Norway | 10 | $1.58 \times 10^{-26}$ | $3.62 \times 10^{-12}$ | 23 |
| Slovenia* | 13 | $7.13 \times 10^{-18}$ | $1.53 \times 10^{-8}$ | 24 |
| Genetic clusters (*K* = 2) |  |  |  |  |
| Cluster 1 | 89 | $2.68 \times 10^{-33}$ | $2.91 \times 10^{-17}$ | 23 |
| Estonia, Latvia, Poland,<br>Russia, Finland, Norway |  |  |  |  |
| Cluster 2 | 30 | $3.54 \times 10^{-22}$ | $3.61 \times 10^{-11}$ | 24 |
| Romania, Slovakia,<br>Croatia*, Slovenia* |  |  |  |  |
| Genetic clusters (*K* = 5) |  |  |  |  |
| Cluster 1 | 47 | $3.01 \times 10^{-33}$ | $5.12 \times 10^{-17}$ | 23 |
| Estonia, Latvia |  |  |  |  |
| Cluster 2 | 15 | $1.11 \times 10^{-28}$ | $1.20 \times 10^{-13}$ | 23 |
| Poland |  |  |  |  |
| Cluster 3 | 17 | $3.76 \times 10^{-34}$ | $1.31 \times 10^{-16}$ | 23 |
| Russia, Finland |  |  |  |  |
| Cluster 4 | 10 | $1.58 \times 10^{-26}$ | $3.62 \times 10^{-12}$ | 23 |
| Norway |  |  |  |  |
| Cluster 5 | 30 | $3.54 \times 10^{-22}$ | $3.61 \times 10^{-11}$ | 24 |
| Romania, Slovakia,<br>Croatia*, Slovenia* |  |  |  |  |

$PID_{unb}$ (unbiased probability of identity) and $PID_{sib}$ (probability of identity given siblings) were calculated using GIMLETv1.3.3 (Valière, 2002)
\* reintroduced population
<sup>a</sup> loci were ranked by informativeness (per sample set), and then the number of loci was determined for which $PID_{sib}$ was $<10^{-4}$ .

**Figure 4.**
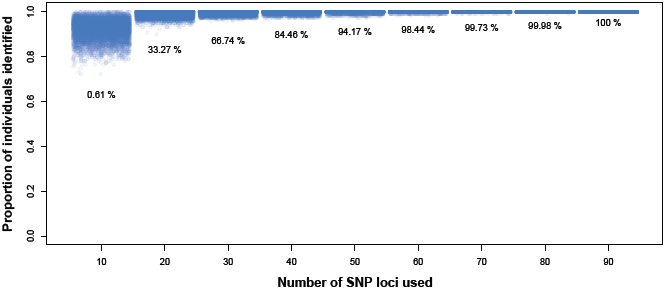
Comparison of the proportion of individuals recovered using different subsets of the SNP data, ranging from 10 to 90 loci. For each subset size 10,000 permutations (random selection of loci without replacement) were plotted; percentage values indicate the number of permutations in which all individuals in the dataset (*N =* 119) were identified. Scatter was added on the x-axis for each subset to visualize density.

### Population structure inferences

The PCA showed a clear separation of two distinct clusters along the first principle component axis (Fig. 5A), which explained 18.1% of variation. These two clusters corresponded to the Eurasian lynx subspecies *Lynx lynx lynx* (left) and *L. l. carpathicus* (right), which can also be differentiated using microsatellites (e.g. Ratkiewicz et al., 2014; Bull et al., 2016). Some substructure within subspecies is also apparent.

**Figure 5.**
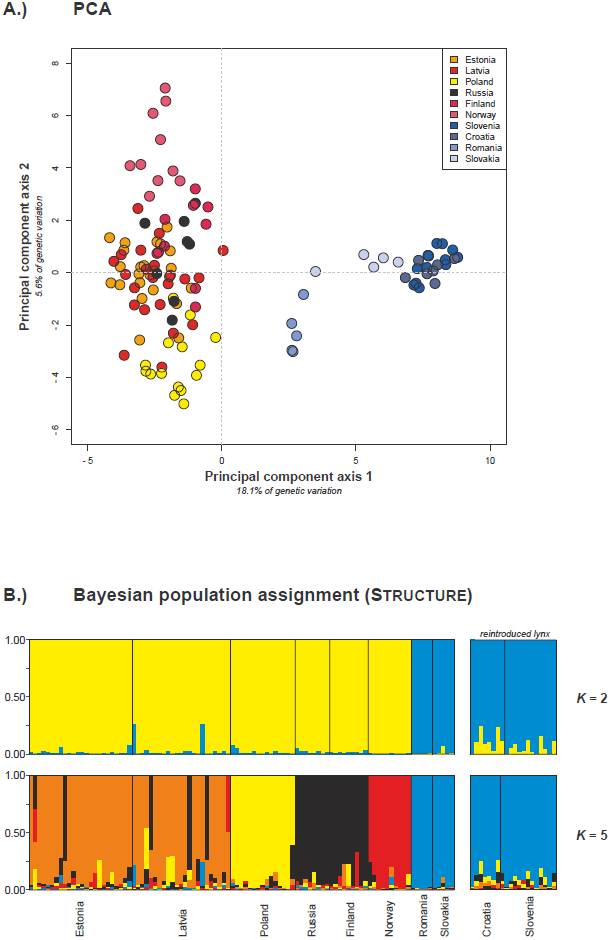
Population structure inferences using the 96 SNP-panel. (A) Principle component analysis (PCA); the first two principle component axes are plotted, with the geographic origin of samples indicated by colour. (B) Bayesian population assignment using STRUCTURE; results for *K =* 2 (top) and *K* = 5 (bottom) are displayed. Results for lynx from reintroduced populations (Croatia, Slovenia) are shown on the right.

In STRUCTURE, the most likely number of genotypic clusters was *K* = 2 (Fig. 5B, upper panel). However, the probability was also high for *K* = 5 (supplemental Fig. S3). Since the Evanno method is designed to detect the highest hierarchical level of genetic structure (Evanno et al., 2005), we also considered *K* = 5 (Fig. 5B lower panel).

Like in the PCA, the two inferred genotypic clusters (*K* = 2) corresponded to the two subspecies *L. l. lynx* (Fig. 5B, upper panel, yellow) and *L. l. carpathicus* (Fig. 5B, upper panel, blue). The lynx from the reintroduced populations in Croatia and Slovenia were assigned with high Q-values (0.75 - 0.99, mean 0.91) to the cluster of their source population (represented by Romanian and Slovakian samples).

The *K* = 5 plot (Fig. 5B, lower panel) displayed a substructure in *L. l. lynx*: individuals from Estonia + Latvia, Poland, Russia + Finland, and Norway now formed their own clusters. There was also admixture between these clusters; most prominently from Russia + Finland into Estonia + Latvia. This extent of detected population genetic structure within *L. l. lynx* is similar to the one previously reported using microsatellites (Ratkiewicz et al., 2014). Again, all lynx from reintroduced populations were assigned with high Q-values (0.73 - 0.98, mean 0.88) to the cluster of their source population.

### Applicability of baits in other taxa

We found high sequence similarity between the cat CDS derived baits and their targets in four other felid species, Iberian lynx, cheetah, leopard and tiger (all four: median sequence divergence of 0.8%; Fig. 6). As expected, sequence divergence between cat CDS derived baits and their targets in other, more distantly related carnivoran species was higher, with a median sequence divergence of 5.8% (Fig. 6).

**Figure 6.**
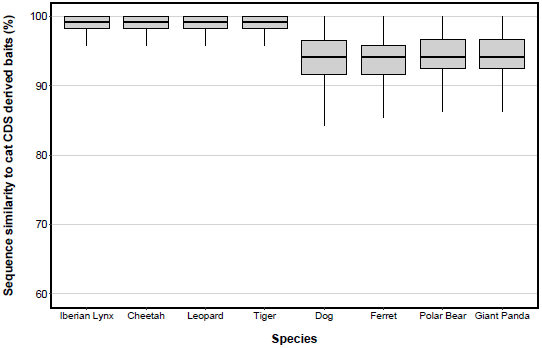
Sequence similarity between the domestic cat CDS derived baits and their targets in other carnivorans with published genomes, including four felids (Iberian lynx, cheetah, leopard, and tiger) and four caniform carnivorans (dog, ferret, polar bear, and giant panda).

## DISCUSSION

Our results show that cross-species capture of coding DNA sequences (CDS) can be used for SNP discovery in non-model organisms, and that a subset of the identified SNPs can be successfully implemented in a high throughput genotyping platform to accurately identify individuals and to infer population genetic structure of the species of interest.

Using publicly available genomic resources for model organisms, we were able to design baits for 100s of target CDS loci that were then enriched in our non-model study species, the Eurasian lynx *Lynx lynx.* We successfully surveyed intraspecific variation in *L. lynx* across its European range, and generated a large data set of SNPs inside CDS and their flanking regions. A large proportion of CDS had good or complete coverage of the target-region (≥15× depth) and yielded 1186 SNP loci for downstream applications.

### Cross-species capture for SNP discovery

Hybridization capture does not require an exact sequence match between bait and target for successful enrichment. While decreasing sequence similarity between bait and target reduces the efficiency of capture (Paijmans et al, 2016), successful enrichment has been reported for species with up to 40% sequence divergence (Li et al., 2013). This ‘mismatch tolerance’ is utilized in cross-species capture, where baits designed using molecular data of one species are used to enrich complementary sequences in one or more other species. In interspecific studies, this yields comparable data (100s-1000s loci) across distantly related species (Lemmon et al., 2012; Faircloth et al., 2012; Bi et al., 2012).

Rather than generating comparable data across multiple species, we used cross-species capture to gain comparable data across many samples within the same species. In this manner, we generated high sequence coverage for a small portion of the nuclear genome – the targeted CDS – across all samples. This portion of the genome was then screened for variation in the form of SNPs, a subset of which was then used to design the SNP-panel.

Hybridization capture is scalable, with the number of targets determined by bait design. To ensure the recovery of sufficient variation for the development of a 96 SNP-panel, we chose to target several hundred CDS spread throughout the genome. However, considering that we far exceeded the number of SNPs required for the development of a 96 SNP-panel, a smaller number of targets would have been sufficient. For the purpose of developing a SNP-panel of similar size, we would still recommend a number of target loci exceeding what is practical using DIY protocols (e.g. 51 loci, Peñalba et al., 2014). Especially, since the per-sample costs of using custom baits (e.g. from MYcroarray) can be reduced by using smaller reaction volumes in combination with a dilution of synthetic RNA-baits (this study; Li et al., 2013; Cruz-Dávalos et al., 2016) and by pooling barcoded libraries of multiple samples prior to hybridization (e.g. Portik et al., 2016; Cruz-Dávalos et al., 2016). In addition, lowering the sequence depth requirement for SNP calling permits pooling of more samples during sequencing and can further reduce costs (e.g. ≥8× depth, Lim & Braun, 2016). Considering our results, and setting a limit of one SNP per target locus for the SNP-panel, we would recommend a minimum of ~250 target loci (400 bp or longer) for SNP discovery using an approach like the one described here.

The increasing availability of genomic resources for non-model organisms, particularly annotated genomes and transcriptomes, improves the chances of finding species for bait-design that are not too distantly related to the target species. While successful enrichment of sequences has been reported for ‘bait species’ with very high divergence times from the target species (up to ~250 million years; Li et al., 2013; Hedtke et al., 2013), several studies examining capture efficiency on a per-target basis (e.g. per exon) have observed a drop in capture efficiency when sequence divergence between bait and target reaches 5-10% (Vallender, 2011; Bi et al., 2012; Peñalba et al., 2014; Bragg et al., 2016). Such a drop in the performance of enrichment would require greater sequencing effort to achieve good sequence coverage of target loci and would thus increase costs for surveying intraspecific variation using cross-species capture. For this reason, we examined the taxonomic breadth at which the baits used here should still perform well. The relatively limited sequence divergence between the cat CDS derived baits and their targets in other carnivorans with published genomes (Iberian lynx, cheetah, leopard, tiger, domestic dog, ferret, polar bear and giant panda; Fig. 6), suggests a relatively broad utility of these baits. This *in silico* assessment showed that the felid species (Iberian lynx, cheetah, leopard and tiger) have low sequence divergence between bait and target, suggesting that the baits we used to enrich CDS in lynx ought to perform very well for other species in the family *Felidae* (consisting of 14 genera and 37 species) that diverged approximately 11 million years ago (Johnson et al. 2006; Li et al., 2016). Unfortunately, except for felids, no other species of the feliform suborder of carnivorans have a published genome (yet); this suborder includes mongooses, meerkats, hyenas, linsangs, civets, and others (6 families without Felidae, with 83 species) that diverged from felids approximately 38 million years ago (Eizirik et al., 2010). Several species of the more distantly related caniform suborder of carnivorans do have a published genome (domestic dog, ferret, polar bear and giant panda). The caniform suborder (9 families, 161 species) diverged from the feliform suborder approximately 59 million years ago (Eizirik et al., 2010). These species appear to be at the limit of what could be considered suitable in terms of cost-effective use of cross-species hybridization for surveying intraspecific variation (using this set of baits). While somewhat crude, this assessment suggests that this single set of baits is suitable to examine intraspecific variation in many species (across taxonomic families or even suborders) over quite substantial divergence times.

Because hybridization capture can be used on samples of poor quality (non-invasively collected material, Perry et al., 2010; archival material, McCormack et al., 2016; Lim & Braun, 2016; ancient material, Carpenter et al., 2013; Enk et al., 2014) it is valuable for studies of taxa for which samples are difficult to obtain for genetic analyses (rare and elusive species, those in difficult to reach habitats, and degraded samples such as archival material). For example, non-invasively collected material can be incorporated in the SNP discovery process to cover portions of a species’ distribution without having fresh tissue samples available. Or, archival and ancient samples can be added to existing data sets, providing information about historical (e.g. extinct) populations (Bi et al., 2014; Lim & Braun, 2016).

### Genetic monitoring using SNPs in CDS

The ability to accurately identify and differentiate individuals is central to population monitoring (Frankham et al., 2010). Thus, molecular markers used in non-invasive genetic monitoring must have sufficient power to differentiate individuals – even closely related individuals – and overcome the analytical difficulties often associated with non-invasively collected sample material, namely DNA extractions of low volume and low concentration that are characterized by varying levels of DNA degradation.

For our SNP-panel, we adopted a SNP typing platform, Fluidigm’s Dynamic Array Chips, which has been successfully used to genotype SNPs in a range of non-invasively collected material, such as individual hair, faecal samples, and urine samples (Nussberger et al., 2014; Kraus et al., 2015). Using this platform, we observed a high genotyping success rate for 80% of samples with very low DNA concentrations (0.5 ng/μL), which we used in lieu of non-invasive samples. Conducting multiple replicates per sample is unproblematic, as only a limited amount of template is required per sample, and the costs of genotyping samples are relatively low (Kraus et al., 2015).

Regarding the identification of individuals, the 96 SNP format performed very well. This is in line with previous studies, which found that 40-100 SNP loci performed similarly well or better than the typical number (10-20) of microsatellite loci used for the purpose of individual identification and kinship analysis (e.g. Tokarska et al., 2009; Hauser et al., 2011; Gärke et al., 2012; Morin et al., 2012; Weinman et al., 2015; Kaiser et al., 2017). Here, we observed that a low number (23 or 24) of informative loci was more than sufficient to distinguish individuals of a given population. Our permutation test showed that even with a substantial number of locus drop-outs (up to 30-40%), we were able to distinguish individuals, indicating that this 96 SNP-panel is robust enough for genotyping poor quality (e.g. non-invasively collected) samples.

The ability to make population structure inferences and to assign individuals to populations is an important component of genetic population monitoring, providing information about animal movements and potential gene flow, the impact of habitat fragmentation, degree of inbreeding, and other population parameters (Frankham et al., 2010). In past years, there has been increasing evidence for the suitability of genic SNPs to ascertain population membership of individuals (Freamo et al., 2011; Defaveri et al., 2013; Oliveira et al., 2015; Zhan et al., 2015; Elbers et al., 2016). This has been examined by itself and also in comparison with both traditionally used microsatellite markers (e.g. Defaveri et al., 2013; Elbers et al., 2016) and non-genic SNPs (e.g. Defaveri et al., 2013; Zhan et al., 2015; Elbers et al., 2016); in all cases, genic SNPs performed equally well or better than alternative markers.

Using our CDS derived SNPs we were able to unambiguously delineate the two Eurasian lynx subspecies in our sample set (*L. l. lynx* and *L. l. carpathicus*). Within *L. l. lynx*, which dominated our sample set, further levels of population structure could be resolved. This structure was congruent with the one detected in a larger sample set (*N* = 298) that had been genotyped at 13 microsatellite loci (Ratkiewicz et al., 2014). Similarly, the extent and direction of introgression detected using our 96 SNP-panel mirrored that observed using microsatellites in the aforementioned study. Lastly, using our 96 SNP-panel we were also able to accurately assign individuals from reintroduced populations to the genetic cluster of their source population. The 96 SNP-panel presented here thus appears more than suitable for the genetic monitoring of Eurasian lynx across their European range considering the higher potential for automation of SNP genotyping, better collaboration possibilities, and cost reduction potential (Kraus et al., 2015).

## CONCLUSION

There is growing evidence for the utility of genic SNPs for the genetic monitoring of populations. As demonstrated here, hybridization capture coupled with high-throughput sequencing is well suited for acquiring information regarding such intraspecific variation, even in cases where study species lack genomic resources. With the increasing availability of genomic resources for non-model species, this kind of approach will become more broadly applicable – even though cross-species capture already shows potential to work across substantial divergence times (e.g. Li et al., 2013; Hedtke et al., 2013; Bragg et al., 2016).

Baits do of course not need to be explicitly designed with the aim of discovering SNPs for genetic monitoring purposes. Thus, baits designed for other purposes (e.g. resolving taxonomic uncertainties, Yuan *et al.,* 2016; identifying regulatory sequences, Yoshihara et al. 2016; identifying adaptive genes, Roffler et al. 2016; investigating loci linked to traits, Springer et al., 2015) can be used to screen samples from related species for intraspecific variation.

## ACKNOWLEDGEMENTS

This research was funded by the Leibniz Association (SAW-2011-SGN-3) and by the Deutscher Akademischer Austauschdienst (A/11/93927). The license for lynx live-trapping and blood sampling in Poland was obtained from the National Ethics Committee for Animal Experiments (no. DB/KKE/PL— 110/2001) and the Local Ethics Committee for Animal Experiments at the Medical University of Białystok, Poland (no. 52/2007). Import of samples from Russia was licensed by CITES permission no 12RU000512.

## AUTHOR CONTRIBUTIONS

D.W.F., R.H.S.K., H.B., R.K., C.N. and J.F. designed the study, K.S., A.P.S., and M.S. coordinated sample collection, D.W.F., J.B., M.A., J.L.A.P. carried out the experiments, D.W.F and D.L. analyzed and interpreted data, and D.W.F. and J.F. wrote the manuscript. All authors edited and approved the final manuscript.

## DATA ACCESSIBILITY

Supplementary Tables S1 and S2 and Figures S1, S2 and S3 can be found online with the study; SNP genotypes, the final 96 SNP-panel assay, as well as an R script are deposited on DRYAD under doi: provided upon acceptance. Illumina sequences are deposited in the NCBI SRA under Accession no.: provided upon acceptance.

## Supporting Information

**Table S1.** Additional lynx samples genotyped using the newly developed 96 SNP-panel.

**Table S2.** Summary of CDS markers.

**Figure S1.** A comparison of the number of unique on-target sequences following one capture versus two consecutive captures, for five samples (see also Table 1). Shown are the number of on-target sequences (%), the number of duplicate sequences (%), and the fold-increase in the number of unique on-target sequences.

**Figure S2.** A ‘LD heatmap’ of pairwise *r*^2^ values, for 92 SNP loci (loci with more than 5% missing data were removed).

**Figure S3.** Plot of Δ*K* values (STRUCTURE analysis) generated by STRUCTUREHARVESTER (Earl & vonHoldt, 2012).

